# Local thermal environment and warming influence supercooling and drive widespread shifts in the metabolome of diapausing *Pieris rapae* butterflies

**DOI:** 10.1101/2020.06.29.178087

**Authors:** Emily E. Mikucki, Brent L. Lockwood

## Abstract

Global climate change has the potential to negatively impact biological systems as organisms are exposed to novel temperature regimes. Increases in annual mean temperature have been accompanied by disproportionate rates of change in temperature across seasons, and winter is the season warming most rapidly. Yet, we know relatively little about how warming will alter the physiology of overwintering organisms. Here, we simulated future warming conditions by comparing diapausing *Pieris rapae* butterfly pupae collected from disparate thermal environments and by exposing *P. rapae* pupae to acute and chronic increases in temperature. First, we compared internal freezing temperatures (supercooling points) of diapausing pupae that were developed in common-garden conditions but whose parents were collected from northern Vermont, USA, or North Carolina, USA. Matching the warmer winter climate of North Carolina, North Carolina pupae had significantly higher supercooling points than Vermont pupae. Next, we measured the effects of acute and chronic warming exposure in Vermont pupae and found that warming induced higher supercooling points. We further characterized the effects of chronic warming by profiling the metabolomes of Vermont pupae via untargeted LC-MS metabolomics. Warming caused significant changes in abundance of hundreds of metabolites across the metabolome. Notably, there were warming-induced shifts in key biochemical pathways, such as pyruvate metabolism, fructose and mannose metabolism, and beta-alanine metabolism, suggesting shifts in energy metabolism and cryoprotection. These results suggest that warming affects various aspects of overwintering physiology in *P. rapae* and may be detrimental depending on the frequency and variation of winter warming events. Future research is needed to ascertain the extent to which the effects of warming are felt among a broader set of populations of *P. rapae*, and among other species, in order to better predict how insects may respond to changes in winter thermal environments.

## INTRODUCTION

Climate change will expose organisms to unpredictable thermal environments to which they may not be adapted through shifts in seasonality (i.e. later onset of winter and/or earlier onset of spring) and the increased frequency of temperature anomalies (Buckley and Kingsolver, 2011; García-Robledo et al., 2016; Sinclair et al., 2016; Somero, 2010; Somero et al., 2017). Mean atmospheric winter temperatures are increasing at a faster rate than any other season (Allen et al., 2019). According to the latest IPCC special report, winter temperatures have shown increased variability with both hotter mean temperatures and a lower frequency of days below freezing (Allen et al., 2019; Hayoe et al., 2018). Thus, it is imperative to characterize how overwintering organisms respond to warming conditions in order to predict how these species will respond to the future climate.

Diapause is an overwintering strategy for temperate insects, and relies on intrinsic physiological mechanisms (Ragland and Keep, 2017; Ragland et al., 2011) that depress metabolic activity, arrest development, and confer cold and stress tolerance (Koštál, 2006; Koštál et al., 2017). A key trait that underlies cold tolerance during diapause is supercooling (Lee, 2010; Sinclair et al., 2015), a freeze avoidance strategy in which body solutions can drop below 0°C without the formation of ice (Somero et al., 2017). Ice formation is detrimental, particularly when it occurs inside cells, due to the volume expansion of freezing aqueous solutions (Somero et al., 2017). Supercooling is achieved by various biochemical mechanisms that work in concert, such as cryoprotectant metabolites that reduce the freezing point via colligative properties (Koštál et al., 2007), ice binding proteins that hinder the formation and spread of ice crystals (Duman, 2015), and cryoprotective dehydration that decreases the potential for body water to freeze by increasing the concentration of cryoprotectant metabolites and ice binding proteins (Walters et al., 2011). For example, one of the lowest supercooling points was reported in larvae of the arctic beetle *Cucujus clavipes*, which can supercool to -58°C via the synthesis and accumulation of high concentrations of glycerol (4-6 mol L^-1^) (Sformo et al., 2010). Vitrification, the conversion of intra- and extracellular water into solid-state vitreous water that does not expand in volume, is yet another mechanism of freeze avoidance that enables *C. clavipes* larvae to survive temperatures down to -100°C (Sformo et al., 2010).

In addition to the intrinsic physiological mechanisms that enable diapausing insects to survive through months of extreme winter conditions, extrinsic factors such as temperature may also influence overwintering. The Arrhenius relationship (i.e., the Q_10_ effect) predicts that increases in temperature will lead to exponential increases in the rates of biochemical reactions (Somero et al., 2017). Thus, if dormant animals rely on cold temperatures as a means to extrinsically regulate and depress the rates of their physiological processes (Geiser, 2004; Hodek and Hodková, 1988; Snapp and Heller, 1981; Storey et al., 2010), warming could lead to increases in biochemical activity that alter development rates (Buckley et al., 2017), deplete energy reserves (Sgolastra et al., 2011; Williams et al., 2012), and hinder cold tolerance (Coleman et al., 2014; Sobek-Swant et al., 2012). Notably, cryoprotectants can be used as fuel sources (Sinclair, 2015; Sinclair and Marshall, 2018; Storey and Storey, 1986), which could directly impact cold tolerance as warming increases energetic demand. Furthermore, insects may be particularly vulnerable to the increased thermal variability (Colinet et al., 2015; Sinclair et al., 2016) that accompanies winter warming (Allen et al., 2018), as even subtle changes in temperature can have adverse effects on diapause development and subsequent spring eclosion success (Lehmann et al., 2018). As further evidence that winter warming could impact overwintering physiology in insects, many studies have reported evidence of local adaptation of cold tolerance that correlates with local thermal environments (Arambourou and Robby, 2015; Bradshaw et al., 2004; Hoffmann et al., 2005; Kukal and Duman, 1989; Nguyen et al., 2019; Preisser et al., 2008; Vrba et al., 2014), suggesting that warmer winters will lead to evolutionary responses in overwintering physiological traits that support cold tolerance and ultimately influence fitness.

While previous work has established connections between the thermal environment and overwintering physiology in various insects, the extent to which changes in temperature influence the molecular underpinnings of overwintering physiology, such as the metabolome, has not been fully explored. Much of the work to uncover the molecular physiological basis of overwintering physiology in insects has employed targeted metabolomics to profile candidate molecules that are known to play a role in overwintering physiology, such as various cryoprotectants (Colinet et al., 2016; Koštál et al., 2007; Koštál et al., 2011; Koštál et al., 2016; Lehmann et al., 2018; Michaud and Denlinger, 2007). These candidate metabolomics studies have contributed important discoveries that enhance our understanding of the molecular basis of diapause and overwintering physiology in insects, but more studies are needed to expand upon this work to include broader surveys that use untargeted metabolomics, such as (Chen et al., 2021; MacMillan et al., 2016; Williams et al., 2014a). In particular, untargeted metabolomics can be employed in order to limit the ascertainment bias that is inherent in candidate metabolite studies (Feder and Walser, 2005) and ultimately provide a more comprehensive picture of complex molecular physiological processes.

To address this gap in knowledge, we investigated thermal effects on supercooling and the metabolome of North American *Pieris rapae* butterflies, which diapause and overwinter in the pupal stage (Richards 1940). *P. rapae*, or the cabbage white butterfly, is a globally abundant insect species that has a broad distribution that spans thermal gradients across five continents (Ryan et al., 2019). Previous work has shown that populations of *P. rapae* from locations with different winter climates have distinct cold tolerance strategies—i.e., freeze avoidance vs. freeze tolerance (Li et al., 2020; Li and Averenskii, 2007; Sømme, 1982). *P. rapae* from North America and Europe are freeze avoidant (Li et al., 2020; Sømme, 1982), whereas *P. rapae* from eastern Siberia are freeze tolerant (Li et al., 2020, Li and Averenskii, 2007). This suggests that the thermal environment influences various aspects of *P. rapae* overwintering physiology, but also highlights the importance of considering ecological relevance when investigating the physiological mechanisms that underlie cold tolerance. Thus, we first sought to confirm the ecological relevance of supercooling in North American *P. rapae* by comparing the supercooling points of diapausing *P. rapae* that were collected from disparate thermal environments in northern Vermont and North Carolina, USA. Because North American *P. rapae* have been shown to be freeze avoidant (Li et al., 2020), we predicted that the ability to supercool would correlate with the local thermal environments in Vermont and North Carolina. We note that supercooling is but one aspect of cold tolerance and may not fully describe fitness or survival in response to cold stress (Lehmann et al., 2018). Nonetheless, supercooling is a critical trait for freeze avoidant insects and, thus, we use it as a proxy for cold tolerance in the present study. Next, we tested whether acute (hours) or chronic (weeks) increases in temperature influence supercooling in Vermont *P. rapae* pupae. Lastly, we conducted untargeted LC-MS metabolomics to characterize the metabolomic profiles of Vermont *P. rapae* pupae that were exposed to chronic warming. We measured metabolomic profiles of Vermont pupae for which we also measured supercooling points, allowing us to correlate metabolomes to supercooling.

Our experiments were designed to assess the relative effects of predicted near-term future increases in temperature on overwintering physiology; the difference in winter temperatures in Vermont and North Carolina, and the degree to which pupae were exposed to increased temperature, approximate the extent of warming that is predicted to occur in Vermont in the next century, if current trends continue (Fig.1). We predicted that (1) pupae would exhibit supercooling points that reflected the thermal environment of their population of origin, (2) higher temperature during diapause would adversely affect supercooling, and (3) warming would induce changes in metabolite abundances across the metabolome that reflect larger shifts in metabolism. Overall, our results suggest that overwintering physiology in diapausing *P. rapae* is strongly influenced by temperature and that winter warming may have important consequences for the ecological physiology of this species.

## MATERIALS AND METHODS

### Adult butterfly collections and maintenance

We collected approximately 40-50 male and female adult *Pieris rapae* butterflies in mid to late September in 2017 at two locations in northwestern Vermont, at least 15 miles apart (44°29’48.52”N, 73°12’20.19”W and 44°17’10.07”N, 73°14’07.11”W). Adult *P. rapae* butterflies were collected from two locations in North Carolina (35°36’19.47”N, 82°20’07.25”W and 35°36’31.57”N, 82°26’31.33”W). After collection, we kept adults in mesh containers (Carolina Biological Supply, 11” diameter × 12” height) with 10 butterflies in each container under common garden conditions of 24°C, 12:12 Light:Dark photoperiod, 55% relative humidity, and with direct access to sunlight. We fed adults a diet of 10% honey solution on a sponge every 24 hours. After 48 hours post-collection, we isolated females in individual mesh containers, and gave them fresh organic kale leaves on which to oviposit. Fertilized eggs were collected every 24 hours, and placed into common garden juvenile rearing conditions. We note our egg collection methods were designed to collect as many eggs as possible over a short period of time from a relatively limited sample of adults, and thus we allowed adults to mate en masse.

### Juvenile stage rearing and diapause induction

*Pieris rapae* diapause in the pupal stage, with the larval stage as the sensitive, or preparative, stage (Richards 1940). To ensure all individuals entered diapause, we subjected all individuals to short-day photoperiods (8L:16D) starting at the embryonic stage. Upon oviposition, eggs were collected and randomly assigned to treatment groups, and placed into plastic containers (35.6cm length x 20.3cm wide x 12.4cm height) in incubators (Percival model DR-36VL) set to 24°C and 55% relative humidity, with approximately 20 eggs in each container. Due to space limitations inside of our incubators, egg clutches were reared in groups of eggs from multiple females. Thus, we did not keep siblings separate from non-siblings and cannot test for the effect of individual female on SCP (see below). We reared larvae on fresh organic kale leaves that were replaced daily.

Upon pupation, roughly 14 days post oviposition, we placed all individuals into petri dishes (60 × 15mm) and in one of three temperature treatments (described below). We note the following constraints of our experimental design: (1) We cannot determine whether among population differences in supercooling point were due to genetic divergence or maternal effects caused by the local environments from which adults were collected, and (2) we cannot account for the parental source of genetic differences observed among individuals within an experimental group, as individuals may or may not have been siblings—multiple female-male pairs were bred in common garden. We also note, however, that adults were collected from outbred natural populations, and F1 offspring were used in the experiment. Thus, there was no inbreeding over generations in the lab, and likely only minimal lab adaptation. Importantly, all individuals in the study possessed unique genomes, regardless of whether they were siblings, due to genetic recombination.

### Warming treatments in diapausing pupae

Trends in winter temperature (December, January, and February) from 1940 to 2020 in Burlington, VT indicate that winters were 3°C warmer, on average, in 2021 than in 1940 (Fig. 1; Least-squares linear regression; daily high temperatures, *R*^2^ = 0.11, *P* = 0.0013, *y* = 0.027*x* – 54.15; daily low temperatures, *R*^2^ = 0.22, *P* = 0.0000065, *y* = 0.046*x* – 101.8). We calculated the average change in temperature from 1940 to 2020 by averaging the slopes in the regression fits of daily high and low temperatures and multiplying by 80 years (0.027°C per year + 0.046°C per year) **÷** 2 × 80 years = 2.9°C). We used these trends to determine ecologically relevant winter warming treatments consisting of a control treatment, chronic warming treatment, and acute warming treatment. We kept control individuals under a temperature regime with a daily fluctuating temperature of 4°C-8°C, which was established from day and nighttime averages in Vermont October representing autumn temperatures when individuals first enter diapause. We kept control individuals under this regime for the entire experiment. To assess population-level differences in diapause physiology, we only subjected individuals from North Carolina to this control treatment, thus, comparing them to the Vermont control treatment. Only pupae from Vermont were exposed to the warming treatments. We kept the chronic warming Vermont individuals under a temperature regime of 7°C-11°C, representing a 3°C increase from the control group, which is indicative of the winter warming pattern described above. We kept the acute warming individuals under the control conditions of daily fluctuating 4°C-8°C but with three, 24-hour warming events of fluctuating 18°C-23°C on days 25, 50 and 75, which reflect record high winter temperatures observed in Vermont. This temperature regime mimics the hottest recorded diurnal and nocturnal temperatures observed in Vermont winters (National Weather Service Forecast Office, https://w2.weather.gov/climate/local_data.php?wfo=btv, accessed February 2020).

**Figure 1.**
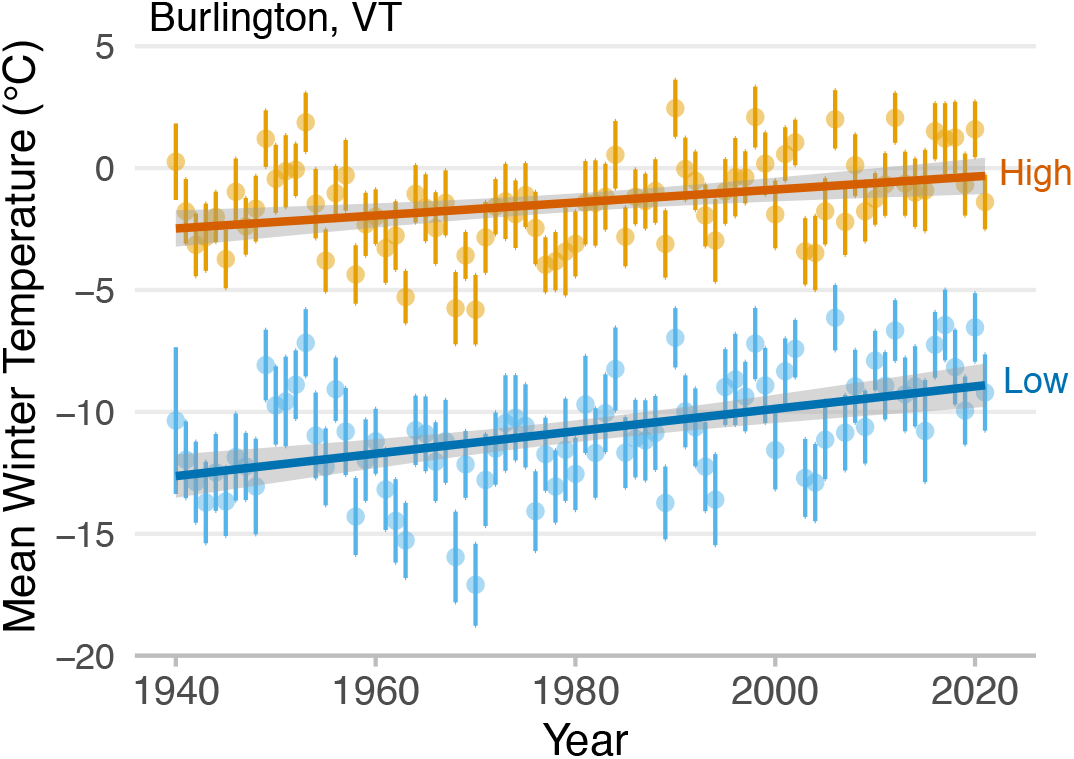
Winter temperature trends in Burlington, VT from 1940 to 2021. Plotted are mean daily high and low temperatures, averaged over December, January, and February for each year. Both daytime highs and lows are warmer now than they were 80 years ago (Least-squares linear regression; High, *R*^2^ = 0.11, *P* = 0.0013, *y* = 0.027*x* – 54.15; Low, *R*^2^ = 0.22, *P* = 0.0000065, *y* = 0.046*x* – 101.8). Error bars and error bands indicate 95% confidence intervals.

### Supercooling point measurement

Diapause length is variable in *P. rapae* and differs by latitude (Saunders, 1982), and we did not measure *in situ* diapause length of the source populations in North Carolina and Vermont. But while pupae in North Carolina are likely to diapause for shorter lengths of time than pupae in Vermont, based on the winter temperatures at both of these sites (Dec., Jan., and Feb.; Figure 2B), we expect that diapause in nature lasts for at least 90 days. We measured the internal freezing temperature, or supercooling point (SCP), of the diapausing pupae based off of the protocols described in Boychuk et al. (2015) and Sinclair et al. (2015) (Boychuk et al., 2015; Sinclair et al., 2015). We weighed individual pupae every 2-4 days using a fine-scale balance at room temperature for no longer than 5 minutes to determine any differences in weight over the course of the 90-day experiment (Mettler Toledo XSE105). We report no statistical differences in weight over time for any of the three treatments (Type II ANOVA, *F*_1,41_=1.62, *P*=0.21). Thus, we did not observe a significant loss of water throughout the experiment. Immediately prior to supercooling point measurement, we weighed each pupa. To determine SCP, we placed pupae attached to a type-K thermocouple wire (OMEGA Engineering) into individual 2-ml microcentrifuge tubes, and then sealed them with parafilm. We equilibrated individuals in a circulating water bath (Polyscience PP15R-30) with Polycool HC -50 anti-freeze liquid, then kept individuals at 0°C for 10 minutes, and then cooled them from 0°C to -30°C at a rate of 0.5°Cmin^-1^. We monitored body temperature using a thermometer and data logger program (OMEGA Engineering HH806AW). SCP was defined as the temperature at which ice formed, and was measured as the lowest temperature (°C) recorded before the detectable presence of an exothermic reaction (ice formation) in the temperature trace. We analyzed individual internal freezing temperatures on Days 25, 50 and 75 in the control (Vermont and North Carolina) and chronic warmed individuals, and 24-hours post-warming in the acute warmed individuals. We measured 4-7 individuals for each temperature by treatment combination. Immediately after SCP analysis, we flash-froze Vermont control and chronic warmed individuals in liquid nitrogen and preserved them at -80°C for metabolomics analysis.

**Figure 2.**
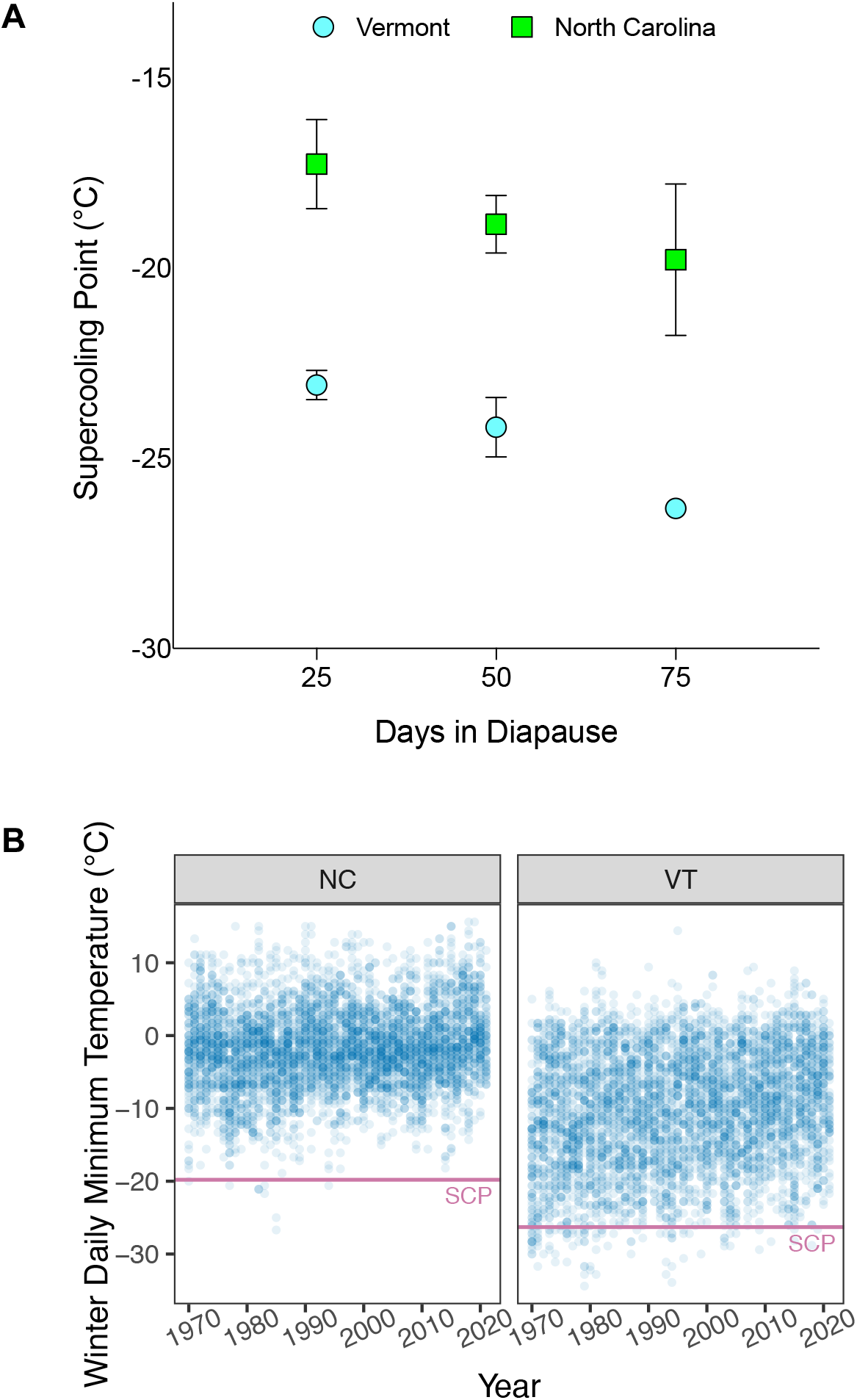
Supercooling points (SCP) of diapausing *Pieris rapae* pupae exposed to control conditions from Vermont and North Carolina populations. (A) SCPs (internal freezing temperatures) were higher in pupae from North Carolina (3-way ANOVA; population factor, *F*_1,24_=60.056, *P*<0.0001; population x day interaction, *F*_1,24_=0.382, *P*=0.543). SCPs decreased through the experiment in both Vermont and North Carolina pupae (3-way ANOVA; day factor, *F*_1,24_=11.466, *P*=0.002). Pupal mass had no effect on supercooling point (3-way ANOVA; mass covariate, *F*_1,24_=0.677, *P*=0.419). Diapausing pupae were exposed to control temperature regime (4-8°C), Vermont (n= 18) and North Carolina (n=14). SCP was measured on days 25, 50, and 75 after diapause induction. SCP is presented as mean freezing temperature (°C) ± standard error of the mean, error bars for control individuals at day 75 are too small to be visible on the plot. (B) Winter daily minimum temperatures (daily lows) in December, January, and February from 1970 to 2021 in Asheville, NC (left panel) and Burlington, VT (right panel). Average supercooling points (SCP) of day 75 pupae collected from NC or VT are indicated by the horizontal pink lines.

We compared supercooling points of (1) Vermont vs. North Carolina populations (under control conditions), (2) Vermont control vs. Vermont chronic warmed, and (3) Vermont control vs. Vermont acute warmed using a linear model incorporating the factors day and treatment as fixed effects and pupal mass as a covariate with a Type II analysis of variance (ANOVA) using the “car” package in R. Pairwise differences between the control and warmed treatments were assessed with Dunnett’s multiple comparison post hoc test using the “lsmeans” package in R. We also investigated trends in winter temperatures at the collection sites in NC and VT and compared them to the observed SCP values for the NC and VT individuals by downloading weather station data from NOAA using the R package “rnoaa” (Chamberlain 2021; NOAA, https://www1.ncdc.noaa.gov/pub/data/ghcn/daily/all, accessed March 2021). The weather station in NC is located at the Asheville Airport, 22 km from the NC collection site (weather station Id: GHCND:USW00003812, 35° 25’ 54.8394”N, 82° 32’ 14.9994”W), and includes a complete daily temperature record from 1964 until the present. The weather station in VT is located at the Burlington Airport, 5 km from the VT collection site (weather station Id: GHCND:USW00014742, 44° 28’ 5.8794”N, 73° 8’ 59.64”W), and includes a complete daily temperature record from 1940 to the present. All supercooling point analyses were performed in R version 4.0.0).

### Global metabolomics sample preparation

We used individual pupae preserved from the supercooling point analysis for global, untargeted metabolomics analysis. To determine the effect of chronic warming on metabolite abundance, we only used control and chronic warmed Vermont individuals (n=17 pupae per group) with at least 4 individuals represented at each timepoint (Days 25, 50 and 75). All samples were sent to the University of Florida Southeast Center for Integrated Metabolomics facility for analysis.

Samples were homogenized in 100 µL 5mM ammonium acetate and protein concentration of each sample homogenate was measured. All samples were normalized to 500 µg/mL protein concentration prior to extraction. Note that because samples were normalized to equal concentrations prior to metabolomics analysis, we did not normalize metabolite abundances to pupal weights. Extraction was performed using protein precipitation. Briefly, 50 µL normalized homogenate was spiked with a mixture of internal standard. Proteins were precipitated by adding 400 µL of 8:1:1 acetonitrile:methanol:acetone. After mixing, proteins were allowed to precipitate for 15 min at 4°C. Supernatant from each sample was collected following centrifugation at 20,000xg for 10 min and dried under a gentle stream of nitrogen at 30°C. Samples were reconstituted with 50 µL of reconstitution solution consisting of injection standards and transferred to LC-vials for analysis.

### LC-MS analysis and data processing

Untargeted metabolomics analysis was performed on a Thermo Q-Exactive Oribtrap mass spectrometer with Dionex UHPLC and autosampler. All samples were analyzed in positive and negative heated electrospray ionization with a mass resolution of 35,000 at m/z 200 as separate injections. Separation was achieved on an ACE 18-pfp 100 × 2.1 mm, 2 µm column with mobile phase A as 0.1% formic acid in water and mobile phase B as acetonitrile. The flow rate was 350 µL/min with a column temperature of 25°C. Injection volume was 2 µL.

MZmine 2.0 was used to identify features, deisotope, align features and perform gap filling to fill in any features that may have been missed in the first alignment algorithm. All adducts and complexes were identified and removed from the data set. This rendered a total of 14,379 features, which we analyzed for significant responses to warming (see below). We used MetaboAnalyst 4.0 (Chong et al., 2019) to normalize the mass spec peak intensities of metabolite features prior to statistical analyses. For each feature, peak intensity was log-transformed and normalized to the sample median. The data were auto-scaled to facilitate comparison among features.

### Statistical analysis of metabolomic data

To test for differences in metabolite abundances between the Vermont control and chronic warmed pupae, we compared the normalized peak intensities, as a proxy for metabolite abundance, of all metabolite features identified by LC-MS. We conducted a principal components analysis to describe the major axes of variation in the dataset, and then tested whether the first principal component (PC1) significantly explained variation in supercooling point among the samples via least-squares linear regression. We then measured the number of metabolites with significantly different peak intensities via Type II ANOVA, with treatment and days in diapause modeled as fixed effects. Features in the positive and negative ion modes were analyzed separately. All p-values were corrected for false discovery via the Benjamini-Hochberg method (Benjamini and Hochberg, 1995). All metabolite features with an FDR < 0.05 were considered to have significantly different abundances. Unless otherwise indicated, we performed all statistical analyses using R version 4.0.0.

### Metabolite annotation and pathway analysis

We used the MS Peaks to Pathways module in MetaboAnalyst 4.0 (Chong et al., 2019) to annotate metabolome features and to conduct pathway analysis. Accurate annotation of untargeted metabolomics data is dependent upon a library of verified standards, which are often incomplete and not representative of the focal species (Li et al., 2013). The MS Peaks to Pathways approach subverts these shortcomings by identifying metabolite sets in the context of KEGG pathways. Metabolite annotation of features is based upon the mass-to-charge ratios in the context of pathways, whose compounds are found to respond in a coordinated manner to experimental manipulation (i.e., warming). Because the goal of this study was to assess the physiological consequences of warming, we focused our pathway analysis and metabolite annotation on the features that were identified to change in abundance in response to chronic warming. We conducted the GSEA algorithm in the MS Peaks to Pathways module of MetaboAnalyst 4.0, which is a rank-based pathway enrichment test. Metabolite features were ranked based on the F-value from the treatment main effect from the ANOVA (see above). We used the *Drosophila melanogaster* KEGG pathway database, which is the only insect species for which KEGG pathway information is available, to identify significantly enriched pathways and metabolites in our dataset. Pathways with an FDR-corrected P-value less than 0.1 were considered significant, following the recommendations of the authors of the analysis software.

## RESULTS

### Effects of population of origin on supercooling

Supercooling point was significantly lower (more negative) in Vermont pupae than North Carolina pupae throughout 75 days of diapause (Fig. 2A). Supercooling point averaged -26.3 ± 0.3°C in Vermont pupae and -19.8 ± 4.0°C in North Carolina pupae at day 75, which corresponds to the disparate extreme low temperatures in these two locations—average extreme minimum temperatures in VT and NC are -29 to -26°C and -18 to -15°C, respectively (Fig. 2B). SCP in both populations decreased through days in diapause.

### Effects of acute and chronic warming on supercooling in Vermont pupae

Chronic and acute warming caused higher supercooling points in Vermont pupae, a pattern that was driven by a marked difference in supercooling between control and warmed pupae at day 50 in diapause (Fig. 2). By day 75, warming had no effect on SCP (Fig. 3). There was an overall decrease in supercooling point over the 75 days in diapause, regardless of treatment, with the lowest average supercooling points on day 75 (Fig. 3).

**Figure 3.**
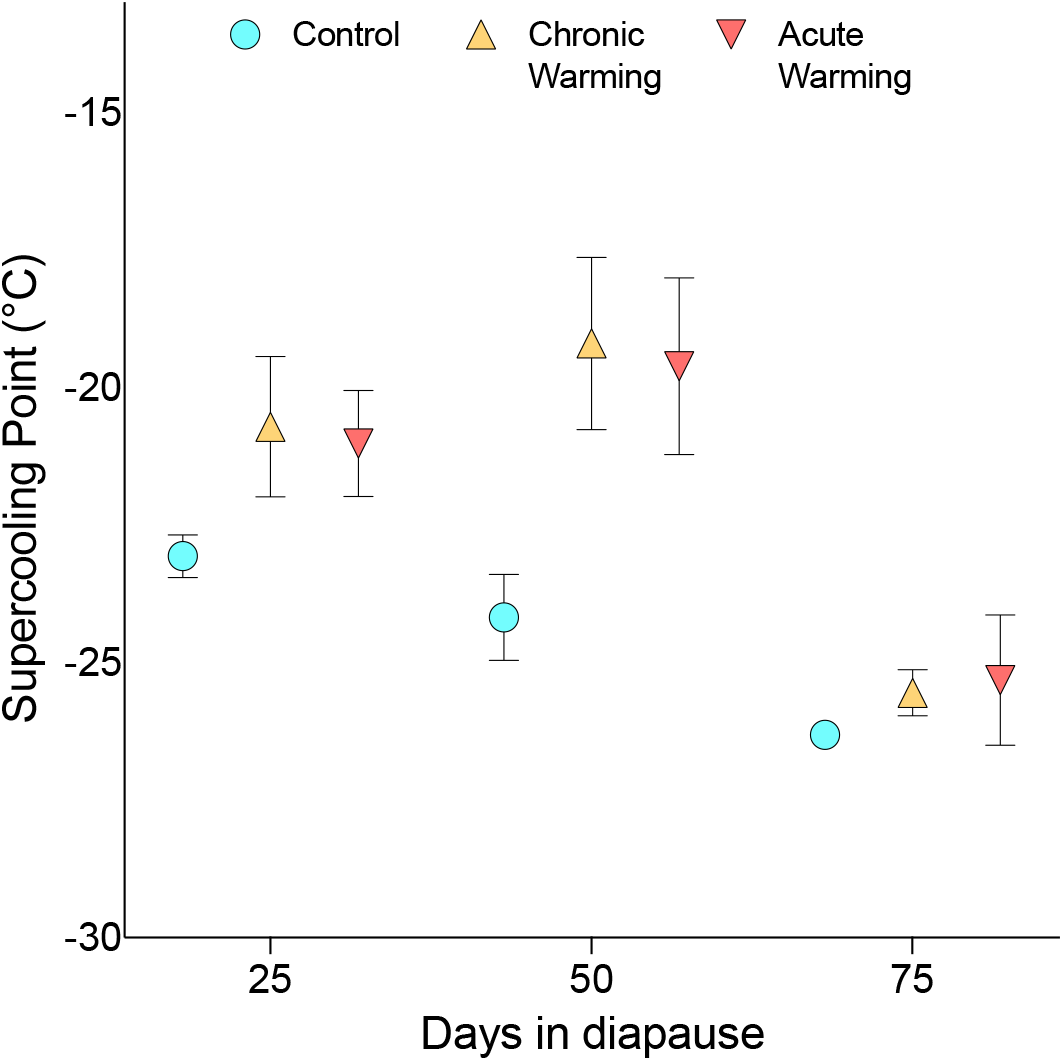
Acute and chronic warming caused higher supercooling points. SCPs (internal freezing temperatures) were higher in pupae exposed to both acute and chronic warming (3-way ANOVA; warming factor, *F*_2,40_=4.928, *P*=0.012). SCP was significantly higher in warmed pupae at day 50 in diapause, but SCP trended higher in warmed pupae even by day 25 (Dunnett’s multiple comparison test, df = 34; day 25 – control vs. acute warming, *t-ratio=*1.545, *P*=0.23, control vs. chronic warming, *t-ratio=*1.582, *P*=0.22; day 50 – control vs. acute warming, *t-ratio=*2.462, *P*=0.036, control vs. chronic warming, *t-ratio=*2.827, *P*=0.015). There was no difference in SCP between control and warmed pupae at day 75 (Dunnett’s test, day 75 – control vs acute warming, *t-ratio=*0.726, *P*=0.69, control vs. chronic warming, *t-ratio=*0.59, *P*=0.81), and SCP decreased overall throughout the experiment (3-way ANOVA, day factor, *F*_1,40_=18.93, *P*<0.0001, warming x day interaction, *F*_2,40_=0.268, *P*=0.77). Mass had no effect on supercooling point (3-way ANOVA, mass covariate, *F*_1,40_=0.829, *P*=0.368). Diapausing pupae were exposed to one of three temperature treatments: control (4-8°C, n=18), acute warming (18-23°C, n=17), or chronic warming (7-11°C, n=17). SCP was measured on days 25, 50, and 75 after diapause induction for the control and chronic warmed pupae, and 24-hr post warming (days 26, 51, and 76) for the acute warmed pupae. SCP is presented as mean freezing temperature (°C) ± standard error of the mean, error bars for control pupae at Day 75 are too small to be visible on the plot.

### Effects of chronic warming on the metabolome in Vermont pupae

Untargeted metabolomics identified a total of 14,379 metabolite features in all pupae from the control and chronically warmed experimental groups. Of these, 1,370 showed significant changes in abundance (normalized peak intensity) through diapause, irrespective of warming treatment (2-way ANOVA, day factor, FDR < 0.01). 443 features showed significant changes in abundance in response to chronic warming (2-way ANOVA, temperature factor, FDR < 0.01), and 16 features showed significant changes in abundance through diapause and in response to warming. No features had abundances that depended on the interaction between day and treatment (2-way ANOVA, day x temperature interaction, all features had an FDR > 0.24).

Metabolite feature abundances representing individual metabolomes revealed that individuals cluster primarily by supercooling point which accounted for nearly 27% of the total variation in abundances of all 14,379 features among pupae (Fig. 4A). In addition, days in diapause accounted for 10% of the total variation in metabolomic profiles (Fig. 4A). The variation among metabolomes, as described by PC1, was strongly correlated to supercooling point (Fig. 4B; Least-squares linear regression of PC1 on SCP, *y = -*15.48*x –* 23.3, *R*^*2*^=0.73, *P*<0.00001).

**Figure 4.**
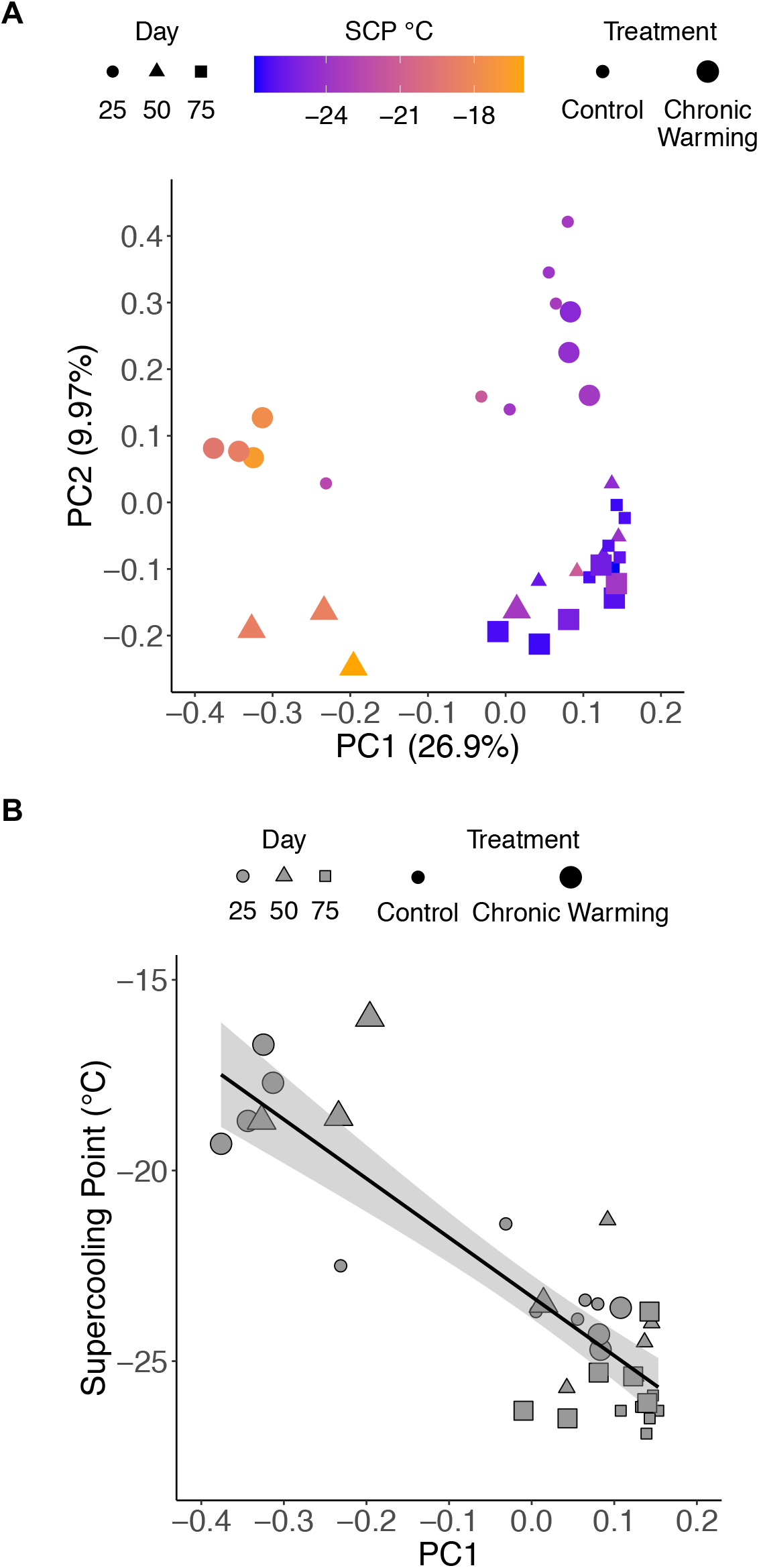
Whole metabolomes cluster by supercooling point (SCP) and days in diapause for control and chronic-warmed Vermont pupae. (A) Principal components analysis of normalized intensities of 14,379 metabolite features among 34 pupae. Each point represents the metabolome of an individual pupa, collapsed in principal component space for the first two principal components that describe 37% of the variation among metabolomes. 26.9% of the variation in metabolomes (PC1) separates pupae by SCP, and 9.97% of the variation (PC2) separates pupae by days in diapause. Day in diapause is indicated by symbol shape, warming treatment is indicated by symbol size, and SCP is indicated by symbol color. (B) Variation among metabolomes is strongly correlated to supercooling point (SCP) (Least-squares linear regression of PC1 on SCP, *y = -*15.48*x –* 23.3, *R*^*2*^=0.73, *P*<0.00001). Day in diapause is indicated by symbol shape and warming treatment is indicated by symbol size for both (A) and (B).

The coordinated changes in the metabolome that accompanied the chronic warming treatment constituted significant changes within 10 biochemical pathways (Fig. 5; Table S1). Chronic warming caused the abundances of metabolites in most (7 out of 10) of these pathways to decrease. Meanwhile, one pathway (arachidonic acid metabolism) showed increases in the abundance of its metabolites. Two pathways (valine, leucine and isoleucine biosynthesis and valine, leucine and isoleucine degradation) showed both increases and decreases in metabolite abundances, and thus these two pathways did not exhibit directionality in warming-induced changes overall (Fig. 5). Three of the pathways—β-alanine metabolism, fructose and mannose metabolism, and glycine, serine and thronine metabolism—implicate the involvement of previously described cryoprotectants, including β-alanine, sorbitol, and glycine (Hahn and Denlinger, 2011; Lee 2010; Michaud and Denlinger, 2007).

**Figure 5:**
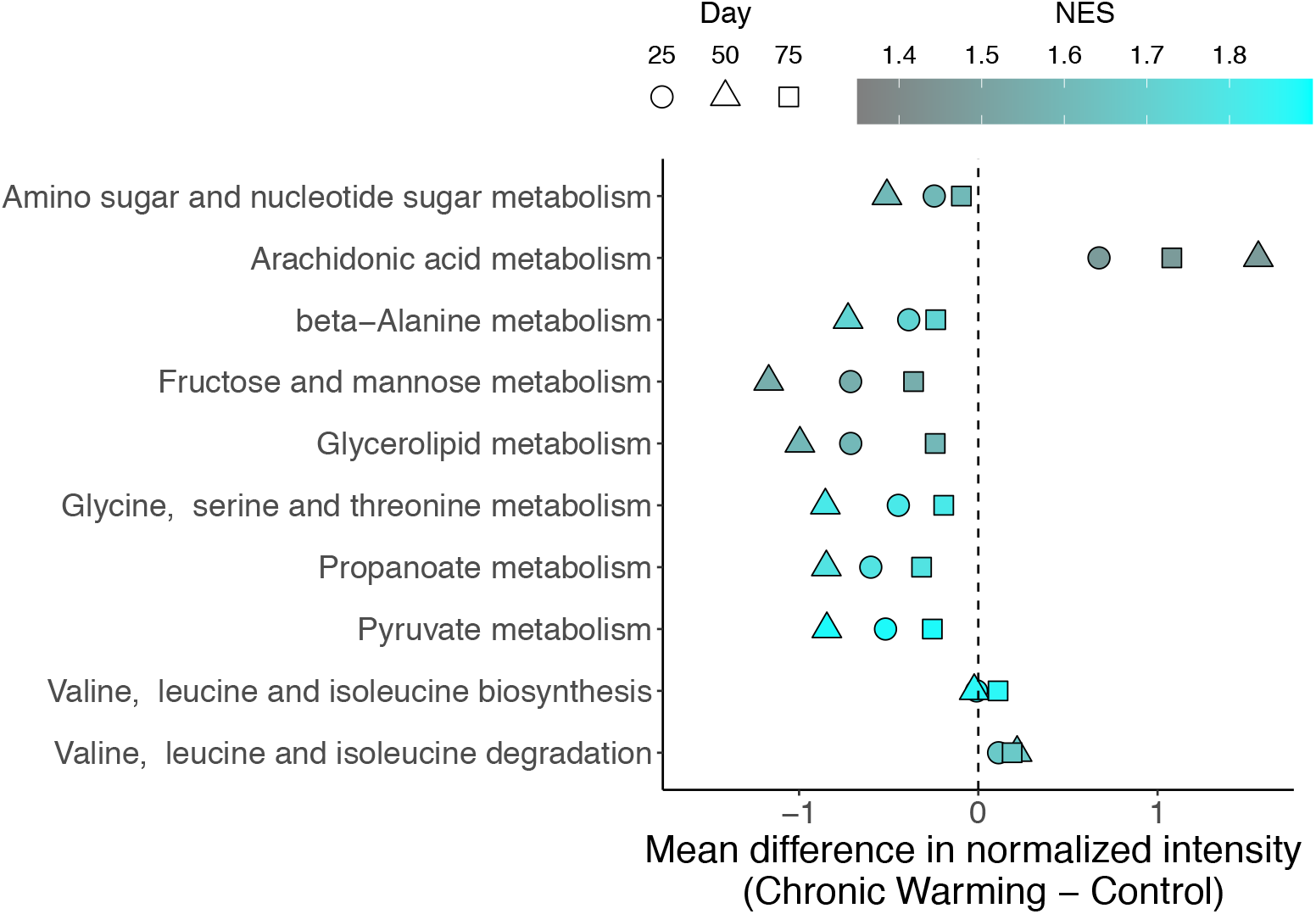
Pathways significantly changed in response to warming. Mean differences in normalized intensity (Chronic warmed – Control), averaged among all features in a given pathway and among pupae in a given day in diapause, for 10 pathways whose member KEGG compounds showed significant differences in normalized intensity in warmed pupae. Represented in the 10 pathways are 153 features that mapped to 81 annotated KEGG compounds. Positive values (x-axis) indicate higher abundances of metabolites in warmed pupae, and negative values indicate lower abundances in warmed pupae, relative to controls. Days in diapause are indicated by the shapes, and normalized enrichment score (NES) is indicated by the color scale. Pathways with higher NES reflect greater proportions of metabolites that were found to be overrepresented in the pathway enrichment analysis. Pathways are listed in alphabetical order.

### Responses of putative cryoprotectants to warming

Pupae with the lowest supercooling points had the highest abundances of three putative cryoprotectants, β-alanine, sorbitol, and glycine, and SCP was negatively correlated with the abundances of all of these metabolites (Fig. 6). Moreover, all three of these metabolites showed significant decreases in abundance after chronic warming (Fig. 6).

**Figure 6:**
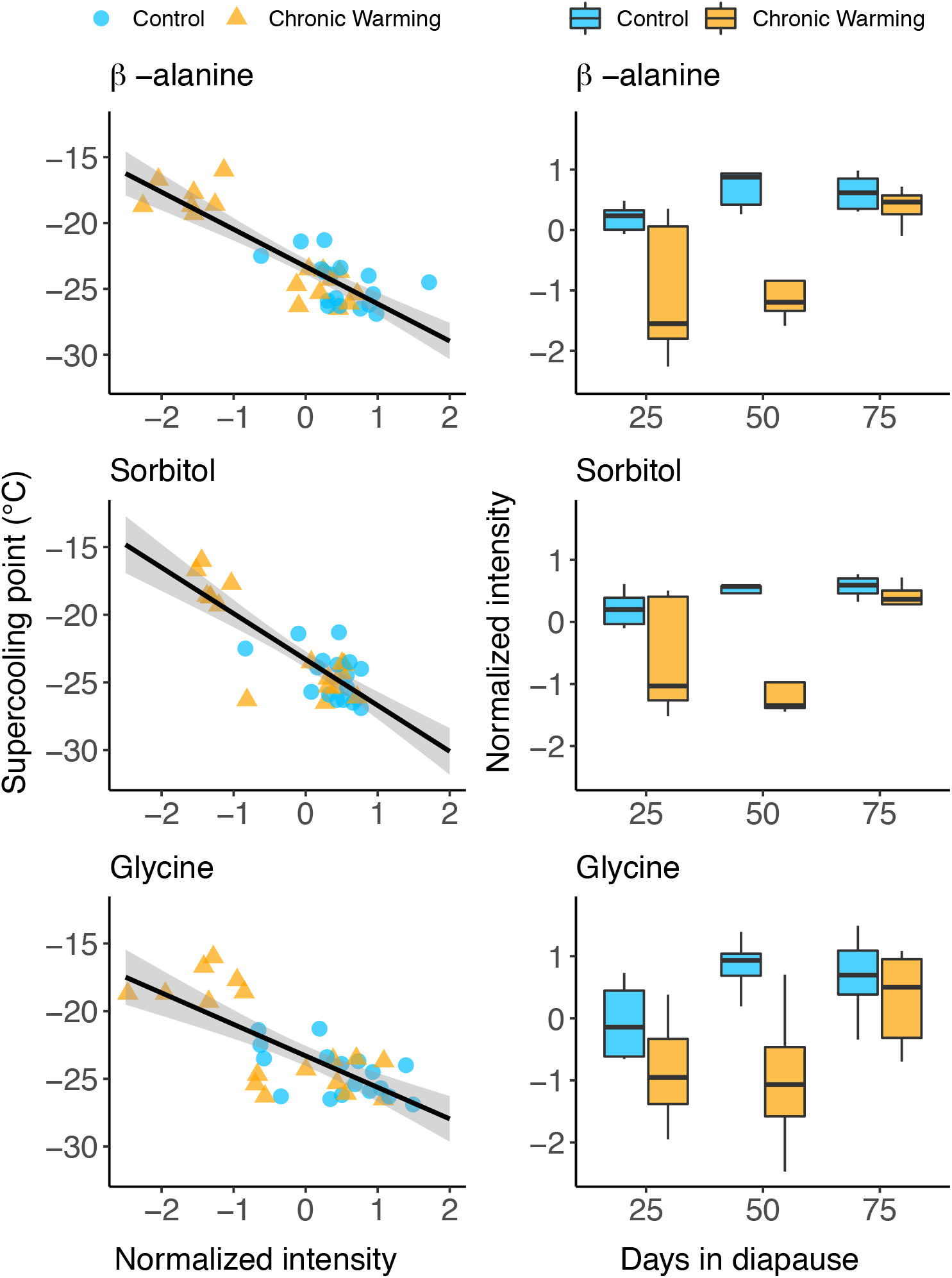
Putative cryoprotectants correlated with SCP and were lower in abundance after warming. (A) β-alanine normalized intensity was negatively correlated with SCP (Least-squares linear regression, *y = -*2.83*x –* 23.31, *R*^*2*^ = 0.72, *P* < 0.00001). (B) β-alanine normalized intensity was reduced after warming (2 way-ANOVA; temperature factor, *F*_1,30_ = 17.16, *P* = 0.0003; day factor, *F*_1,30_ = 12.00, *P* = 0.002; temperature x day interaction, *F*_1,30_ = 2.34, *P* = 0.14). (C) Sorbitol normalized intensity was negatively correlated with SCP (Least-squares linear regression, *y = -*3.4*x –* 23.31, *R*^*2*^ = 0.70, *P* < 0.00001). (D) Sorbitol normalized intensity was reduced after warming (2 way-ANOVA; temperature factor, *F*_1,30_ = 11.11, *P* = 0.002; day factor, *F*_1,30_ = 5.84, *P* = 0.02; temperature x day interaction, *F*_1,30_ = 0.10, *P* = 0.62). (E) Glycine normalized intensity was negatively correlated with SCP (Least-squares linear regression, *y = -* 2.33*x –* 23.31, *R*^*2*^ = 0.55, *P* < 0.00001). (F) Glycine normalized intensity was reduced after warming (2 way-ANOVA; temperature factor, *F*_1,30_ = 11.36, *P* = 0.002; day factor, *F*_1,30_ = 4.52, *P* = 0.02; temperature x day interaction, *F*_1,30_ = 2.30, *P* = 0.12). Data represent mean normalized intensity of all features that matched a given metabolite (β-alanine: 3 features, sorbitol: 11 features, glycine: 1 feature).

## DISCUSSION

A key question in the fields of ecological and evolutionary physiology is how populations and species will respond to future climate change. Our data shed light on this question in two ways. First, we report variation in supercooling among populations of *P. rapae* that matches the historical winter thermal environments in Vermont and North Carolina, USA. Whether or not this difference in supercooling ability is a fixed divergence that resulted from local adaptation is an open question. But regardless of the cause, these data suggest that the natural thermal environment influences overwintering physiology in *P. rapae*, which could have consequences for this species in the future. If there is standing genetic variation in supercooling among populations, then natural selection could act upon this variation and lead to evolutionary responses in supercooling to future winter warming conditions. Second, we demonstrate that warming modified the intrinsic physiological mechanisms that underlie freeze avoidance in diapausing *Pieris rapae* pupae, as warming led to less negative supercooling points that were accompanied by shifts in the metabolomic signatures of cryoprotection and other metabolic pathways. Because supercooling is a key trait for the survival of freeze avoidant overwintering insects, our results suggest that warming could threaten *P. rapae* by raising their supercooling points.

While the present study cannot attribute differences in supercooling between populations of *P. rapae* to a genetic basis, mounting evidence suggests that there is standing genetic variation in various cold tolerance traits in insect populations. In *Drosophila melanogaster*, there is evidence of clinal variation in chill coma recovery time (Hoffmann et al., 2003), and this trait has been shown to evolve in response to laboratory selection (Williams et al., 2014a). In addition, there is a substantial degree of standing genetic variation in whole-organism cold tolerance even in a single population of *D. melanogaster* (Teets and Hahn, 2018). In the pitcher plant mosquito, *Wyeomyia smithii*, classic work by Bradshaw and colleagues (Bradshaw et al., 2004) used reciprocal transplants to demonstrate significant local adaptation in cold tolerance between northern and southern populations. In *Colias* spp. of butterflies, there is evidence of local adaptation in supercooling across elevational gradients that corresponds to winter climates in lowland vs. mountain habitats in Europe (Vrba et al., 2014). Because cold tolerance has evolved in the past, these patterns suggest that overwintering traits could evolve in response to future winter warming; however, the presence of genetic variation in overwintering traits does not necessarily mean that these traits will evolve, even if natural selection is predicted to act upon these traits in a warmer world. It could be that maintaining robust cold tolerance traits, such as supercooling, in itself does not pose a significant cost. Indeed, warmer winters may lead to selection on photoperiodic cues that shorten diapause, rather than direct selection on cold tolerance *per se* (Bradshaw and Holzapfel, 2010; Bradshaw et al., 2004). Ultimately, responses of insect populations to future winter warming will depend not only on the degree of warming but also on the level of thermal variation. Our data suggest that even short-term (24 h) warming may compromise supercooling, which could have deleterious consequences if warm anomalies are followed by extreme low temperatures (see below).

Our results suggest that warming could threaten the survival of *P. rapae* pupae in nature by comprising supercooling ability. Although the observed supercooling points in warmed pupae were relatively low, these supercooling points are within the range of winter temperatures in Vermont. Thus, the warmed pupae from this study could have frozen to death in the wild. Other studies have also shown that increased thermal variability leads to decreased cold hardiness in diapausing insects, including the cabbage root fly, *Delia radicum* (Košťál and šimek, 1995), the emerald ash borer, *Agrilus planipenis* (Sobek-Swant, 2012), the hemlock looper, *Lambdina fiscellaria* (Rochefort et al., 2011; Vallières et al., 2015), and the anise swallowtail butterfly, *Papilio zelicaon* (Williams et al., 2014b).

The timing of winter warming, and subsequent cold challenges, may be an important factor that determines the relative risk of *P. rapae* in warmer winters. Supercooling points decreased throughout diapause in the present study, and many insect species have been shown to follow this same trajectory through diapause (Bale, 2002; Hodek and Hodková, 1988; Marshall and Sinclair, 2015; Pullin et al., 1991), which mirrors decreasing winter temperatures from December through February. Meanwhile, in the present study, pupae at day 50 in diapause showed the largest increase in supercooling points after warming treatment, yet day 75 pupae maintained low supercooling points in spite of warming. The disparate responses of pupae to warming at day 50 vs. day 75 could have been due to increased vulnerability to thermal challenge at day 50 in diapause. Indeed, some stages of diapause may be more thermally responsive than others, as has been observed during metamorphosis in *D. melanogaster* (Merkey et al., 2011). Alternatively, it may not have been the vulnerability of day 50 pupae, but rather the ability of pupae to physiologically acclimate to chronic warming by day 75, such that they could regain low supercooling points. If this were the case, it would help explain the similarity in metabolomes of all day 75 pupae, regardless of warming treatment (Fig. 3A). Thermal acclimation to periodic warming has been previously shown to improve cold hardiness in locust eggs, *Locusta migratoria* (Wang et al., 2006), and adult flies, *D. melanogaster* (Colinet et al., 2016).

We note that extrapolating our results to what pupae experience in nature assumes the direct exposure of diapausing pupae to changes in atmospheric temperatures, which may or may not be a realistic assumption, depending on snow cover that could insulate insects against thermal fluctuations (Boychuk et al., 2015; Sinclair, 2001). However, winter warming is predicted to lead to loss or reduction in snow cover, which would subsequently expose diapausing individuals to fluctuating temperatures and to a higher number of freeze-thaw events (Bale and Hayward, 2010). We also acknowledge that supercooling is a mechanism of cold tolerance that is most critical for species that are freeze avoidant—i.e., species that cannot survive if they experience internal freezing. Supercooling may or may not be important for overwintering survival in species that are freeze tolerant—i.e., species that can survive internal freezing (Marshall and Sinclair, 2012). It remains to be determined whether freeze tolerant species will be challenged by winter warming; however, thermal acclimation influences cold tolerance in at least some freeze-tolerant insects (Li et al., 2020; Toxopeus et al., 2019), suggesting that winter warming may also impact species that are freeze tolerant and not only species that are freeze avoidant, as we have shown here.

Our untargeted metabolomic analysis identified more that 14,000 metabolite features, and the results reveal three major findings. First, supercooling in *P. rapae* diapausing pupae may be influenced by the abundances of thousands of metabolites, as a large proportion of the variance (27%) in abundances of metabolites across the whole metabolome and among pupae significantly correlated to supercooling point. Previous work has established a solid paradigm for interpreting the relationship between cold tolerance traits and metabolite abundance.

Overwintering insects accumulate higher concentrations of key metabolites, or cryoprotectants, to lower the freezing point of intra- and extra-cellular solutions (Bale, 2002; Storey and Storey, 1988; Storey and Storey, 1990). Thus, it is perhaps not surprising to see correlations between metabolite abundance and supercooling, particularly among putative cryoprotectants, such as β-alanine, sorbitol, and glycine (Michaud and Denlinger, 2007; Michaud et al., 2008; Storey and Storey, 1990). In addition, sorbitol and glycine have been shown to stabilize macromolecular structures like proteins (Street et al., 2006; Yancey et al., 1982), suggesting other potential benefits of these compounds in addition to freezing point depression. Targeted metabolomics studies that measure tens-to-hundreds of metabolites corroborate the relationship between metabolite abundance and various cold tolerance traits, including supercooling (Koštál et al., 2007; Koštál et al., 2011; Lehmann et al., 2018; Michaud and Denlinger, 2007). Our data provide a broadscale perspective on the potential role of the metabolome in setting lower thermal limits, which extends previous work to implicate the involvement of a wide array of molecular players.

Alternatively, warming-induced changes in supercooling point and metabolite abundances, including cryoprotectants, could have occurred independently from one another. It is important to note that the present study does not fully account for the diversity of potential mechanisms that underlie supercooling. A critical factor that influences the supercooling point is osmolality, which we did not measure in this study. Dissolved solutes in an insect’s hemolymph colligatively lower their freezing point by 1.86°C per osmole of solute (Denlinger and Lee, 2010), but other non-colligative mechanisms, such as ice-binding proteins, can contribute to the lowering of supercooling point as well (Meister et al., 2013). If non-colligative mechanisms contribute to supercooling in diapausing *P. rapae* pupae, then the metabolomic data we present herein would not fully describe the mechanisms that underlie the shifts in supercooling point that occurred in response to warming. Nonetheless, the correlation between the metabolome and supercooling is noteworthy and deserves further investigation. Future studies should measure osmolality (indicative of colligative mechanisms) and thermal hysteresis (indicative of non-colligative mechanisms) in response to warming to provide further insights into the connection between putative cryoprotectant metabolites and supercooling in overwintering species.

The second major finding of the metabolomics screen is that chronic warming (+3°C) caused shifts in core metabolic pathways, suggesting that even subtle changes in temperature cause changes in metabolism during diapause. Overwhelmingly, warming caused metabolite abundances to decrease; for example, metabolites within the fructose and mannose metabolism pathway, glycerolipid metabolism pathway, and pyruvate metabolism pathway were all significantly higher in control individuals (Fig. 3). Previous work has shown that metabolomes are dynamic, shift throughout diapause, and respond to temperature (Koštál et al., 2007; Koštál et al., 2011; Lehmann et al., 2018; Michaud and Denlinger, 2007). We noted shifts in the metabolome through time, as the second main axis (PC 2), which accounted for approx. 10% of the variation in metabolomic profiles among pupae, separated day 25 pupae from day 50 and day 75 pupae. But regardless of these metabolomic shifts that occurred through diapause, many of the changes in metabolomic profiles were induced by warming, particularly at day 50 in diapause. Moreover, many of the pathways that shifted in response to warming are involved in energy metabolism, such as pyruvate metabolism. Warming-induced decreases in the metabolites involved in pyruvate metabolism could indicate alterations in glycolysis or glycogenolysis, suggesting that ATP generating pathways could respond to winter warming exposure (Denlinger and Lee, 2010). This is of particular concern because diapausing pupae should be metabolically quiescent and able to maintain stable metabolism throughout diapause. Yet, if warming increases energy metabolism, then this could lead diapausing pupae to deplete their energy reserves. In addition, the maintenance of cold tolerance during diapause is dependent on the availability of energy reserves, as fuel sources (lipids, carbohydrates, and amino acids) are also used as anti-freezing cryoprotectants (Denlinger, 2002; Hahn and Denlinger, 2011; Storey and Storey, 2012). We did not assay total lipid or sugar content, nor did we measure metabolic rates; thus, the significance of these findings remains unresolved. Nevertheless, the characterization of energetics in the context of warming in diapause is likely to be a worthwhile avenue of future research.

Third, untargeted metabolomics elucidated patterns of metabolite abundances that we would not have otherwise seen if we had taken a targeted metabolomics approach. For example, the arachidonic acid metabolism pathway was the only pathway in which metabolites exhibited increased abundances in warmed pupae (Fig. 3). The specific function of arachidonic acid in diapausing *P. rapae* pupae remains to be determined, but based on what is known about the role of arachidonic acid in hibernating mammals, this result suggests that winter warming may impact the utilization of energy stores. Arachidonic acid is a long-chain fatty acid that has been shown to regulate the activity of peroxisome proliferator-activated receptor α, a protein involved in mobilizing lipid stores (Wu et al., 2001) and a key regulator of lipid metabolism upon entrance into hibernation in ground squirrels, *Spermophilus tridecemlineatus* (Buck et al., 2002).

Functional data on arachidonic acid in insects is lacking; thus, the potential role of arachidonic acid in regulating energetic processes during diapause remains obscure. It has been shown that arachidonic acid is a key polyunsaturated fatty acid in the cellular membrane phospholipids of *Manduca sexta* (Ogg et al., 1991) and arachidonic acid is down-regulated in diapausing pupae of the flesh fly (*Sarcophaga crassipalpis*) following acute cold stress (Michaud and Denlinger, 2006). But future study is needed to unravel the potential role of arachidonic acid in the context of diapause and environmental change in insects.

### Winter warming: good or bad?

Whether winter warming will benefit or hinder overwintering organisms is currently under debate. Some research argues that warmer winter temperatures will result in beneficial effects on temperate species, as these warmer patterns could lead to increased survival, decreased cold-induced stress, and the ability to expand geographic ranges (Crozier, 2003). If cold stress lowers survival in ectothermic organisms, then the predicted 1°C-5°C increase in winter temperatures could increase survival through winter (Bale and Hayward, 2010). Although this prediction may be true for some species, based upon the data we present herein, not all overwintering organisms will benefit from winter warming. Indeed, previous research on *Pieris napi* has shown that chilling and cold temperatures are needed for endogenous diapause to maintain its developmental trajectory and to progress to post-diapause quiescence for spring emergence (Lehmann et al., 2018; Posledovich et al., 2015). Thus, warming could disrupt the transition into post-diapause development, leading to a longer diapause state or decreased eclosion success. An additional consequence of winter warming may be the earlier spring emergence of insects, including many butterfly and bee species (Bartomeus et al., 2011; Bosch and Kemp, 2003). This phenomenon has potentially negative effects if it causes asynchrony with insects’ host plants or if individuals experience severe environmental conditions post-emergence. Warming may also lead to shifts in the diapause program, including delays to diapause entry, decoupling of environmental cues (temperature and photoperiod) that maintain diapause, and/or the elimination of diapause completely (Bale and Hayward, 2010; Hodek and Hodková, 1988). Thus, at least for insects that have evolved to overwinter in a dormant state, winter warming may pose a significant environmental challenge (Stuhldreher et al., 2014), despite the presence of conditions that are seemingly less harsh.

## CONCLUSION

Our study provides a molecular physiological perspective on the effects of temperature on the physiology of overwintering insects, thus providing insights into the challenges that species may endure as winter temperatures increase and fluctuate with climate change. Future research exploring the effects of warming on overwintering organisms should address not only the direct effects of warming on physiological mechanisms and maintenance, as measured here, but also pre- and post-winter development and subsequent reproductive success after eclosion. Furthermore, research should also focus on comparing populations, ideally in a reciprocal transplant experimental design, to better understand population-level responses and assess the degree to which overwintering traits are locally adapted. This will allow us to better predict the adaptive potential of overwintering traits in the face of winter warming.

## Acknowledgments

We thank Caitlin Decara, Hannah Lewis, and Grace Seta for butterfly collections and larval stage husbandry. We thank Amy Boyd and her students at Warren Wilson College for North Carolina butterfly collections. We thank the University of Florida Southeast Center for Integrated Metabolomics facility for processing the metabolomics samples. We thank Alison Brody, Ingi Agnarsson, Sara Helms Cahan, Sumaetee Tangwancharoen, Thomas O’Leary, and the two anonymous reviewers for helpful comments on this manuscript. This work was supported by National Science Foundation grant IOS-1750322 to B.L.L, and the John Wheeler and Dr. Roberto Fabri Fialho Endowments through the University of Vermont to E.E.M.

## Author contributions

E.E.M conducted the experiment. E.E.M and B.L.L performed data analysis and wrote the manuscript.

## Supplemental Tables

**Table S1.** KEGG pathways significantly enriched in the metabolomic response to warming (MS Peaks to Pathways, MetaboAnalyst)(Chong et al. 2019).

| Pathway | Total Compounds | Compounds in Dataset | *P-val* | Adjusted *P-val* | NES |
| --- | --- | --- | --- | --- | --- |
| Glycine, serine and threonine metabolism | 25 | 17 | 0.010 | 0.079 | 1.81 |
| Amino sugar and nucleotide sugar metabolism | 34 | 20 | 0.010 | 0.079 | 1.607 |
| Pyruvate metabolism | 24 | 15 | 0.010 | 0.079 | 1.885 |
| Fructose and mannose metabolism | 18 | 13 | 0.010 | 0.079 | 1.562 |
| beta-Alanine metabolism | 13 | 8 | 0.011 | 0.079 | 1.724 |
| Valine, leucine and isoleucine degradation | 35 | 10 | 0.011 | 0.079 | 1.676 |
| Valine, leucine and isoleucine biosynthesis | 13 | 9 | 0.011 | 0.079 | 1.869 |
| Propanoate metabolism | 18 | 9 | 0.011 | 0.079 | 1.786 |
| Glycerolipid metabolism | 16 | 6 | 0.011 | 0.079 | 1.604 |
| Arachidonic acid metabolism | 10 | 2 | 0.014 | 0.087 | 1.473 |

